# Tau avoids the GTP cap at growing microtubule plus ends

**DOI:** 10.1101/2019.12.31.891234

**Authors:** Brian T. Castle, Kristen M. McKibben, Elizabeth Rhoades, David J. Odde

**Author notes:** Correspondence to David J. Odde.

## Abstract

Plus-end tracking proteins (+TIPs) are a group of proteins that associate with the growing end of microtubules and mediate important cellular functions including neural development and cell division. Work in recent years has shown that the majority of +TIPs are directed to the plus-end through a family of end binding proteins (EBs), which preferentially bind the stabilizing cap of GTP-tubulin present during microtubule growth, versus weaker binding to GDP-tubulin in the proximal microtubule. One question yet to be addressed is whether there may exist other microtubule associated proteins (MAPs) that preferentially bind specific nucleotide states of tubulin. Here we report that the neuronal MAP tau, which is enriched in axons where it promotes microtubule growth and bundling, preferentially binds GDP-tubulin (*K*_D_ = 0.26 µM) over GMPCPP-tubulin (*K*_D_ = 1.1 µM) *in vitro* as well as GTP-tubulin at the tips of growing microtubules *in vitro* and *in vivo*. This nucleotide preference causes tau binding to lag behind the growing microtubule plus-end by about 100-200 nm both *in vitro* and in live cells. Thus, tau is a microtubule tip avoiding protein, establishing a new class of tip avoiding MAPs, and acts primarily by suppressing microtubule shortening rather than promoting growth. We speculate that neurological disease-relevant tau mutations may exert their phenotype by their failure to properly recognize GDP-tubulin, thus displacing +TIPs, such as EB3, and their associated activities into abnormal locations in the neuron.

## Introduction

Microtubules serve essential roles in cellular processes such as cell division and neuronal development. To serve their function, microtubules rely on a dynamic assembly process known as dynamic instability^1^, where microtubules switch stochastically between alternate phases of net growth and net shortening. This switching is tightly coupled to the presence or absence of a stabilizing region of GTP-bound tubulin at the microtubule end^2^. When the stabilizing region is lost, unstable GDP-bound tubulin is exposed and the microtubule rapidly disassembles until a stabilizing region of GTP-tubulin is re-established. This dynamic switching is regulated by an array of microtubule-associated proteins (MAPs), which influence one or more aspects of dynamic instability as well as the interaction between microtubules and other intracellular structures.

One group of MAPs in particular, known as plus end tracking proteins (+TIPS), preferentially binds the growing microtubule end over those that are shortening. Recent work has shown that many +TIPs localize to the growing end through an interaction with end binding protein 1 (EB1) (reviewed in ^3^), which is believed to preferentially bind the stabilizing region of GTP-tubulin at the microtubule plus end^4, 5^. While much research has focused on characterizing the binding of +TIPs, much less has focused on potential preferential binding of MAPs that is not specific to the microtubule plus-end. One explanation could be that those preferentially binding to GDP-tubulin would bind along the length of the microtubule lattice, and not be readily detectable compared to the signature comet of EBs and other +TIPs localized solely to the end of microtubules. Some evidence has suggested that tau, a neuron-specific MAP, has the ability to recognize different nucleotides states^6, 7^. Although, these studies either examined a low concentration of tau^7^, below the concentrations at which it has been shown to influence microtubule dynamics^8, 9^, or examined tau binding to microtubules stabilized by paclitaxel^6^, which influences tau’s interaction with microtubules^10^. Therefore, a more complete examination of tau’s potential tubulin nucleotide binding preference is needed with intact microtubules in the absence of drugs that influence microtubule assembly.

To better understand tau’s interaction with microtubles, we examine the ability of the longest isoform of human tau, 2N4R, to recognize different nucleotide states of tubulin, stable and unstable, both with purified protein *in vitro* and in mammalian cells using fluorescence microscopy. We find that tau preferentially binds less stable (GDP) over more stable (GTP or GMPCPP) regions of the microtubule *in vitro*. This results in a paucity of tau at the microtubule plus end during periods of growth, but not during shortening, both *in vitro* and *in vivo*, the exact opposite of a +TIP protein. Furthermore, we find that the main effect of tau is to inhibit shortening and rescue *in vivo*, consistent with its avoidance of the growing plus end. We further show that these results are most compatible with a nucleotide preference model and cannot be explained simply by tau binding kinetics or a model in which tau competes with other MAPs for binding. Thus, tau is a microtubule tip avoider, preferentially binding GDP-tubulin to stabilize the microtubule against shortening, thereby establishing the ability of MAPs to avoid GTP-tubulin at the tips of growing microtubules.

## Results

### 2N4R tau preferentially binds GDP microtubules in vitro

Recent evidence suggests a potential for tau to recognize different nucleotide states^6, 7^. These studies, however, either examined binding at a single tau concentration (10 nM), well below that shown to influence microtubule assembly dynamics^8, 9^, or used paclitaxel-stabilized microtubules in solution. Here we sought to assess tau’s ability to recognize nucleotide states directly, in the absence of any potential influence of taxol^11^. To determine if tau preferentially binds to different tubulin nucleotide states we measured the number of Alexa488 labeled tau molecules (Alexa488-tau) bound to different regions of microtubules grown from microtubule seeds stabilized with GMPCPP, a non-hydrolyzable analog of GTP. With 1 mM GTP present in the imaging solution, a stabilizing cap of GTP-tubulin is hydrolyzed to GDP-tubulin as the microtubule assembles off the seed. Therefore, the majority of the microtubule extension will be GDP-tubulin. In order to identify the different regions, microtubule seeds were labeled with 15% rhodamine-tubulin while only 5% rhodamine-tubulin was added to the imaging solution (Figure 1A). As seen in Figure 1B, tau is enriched on the GDP-tubulin extensions compared to the GMPCPP seeds. This preferential labeling was not a function of the rhodamine labeling, as the same result was seen when the labeling ratio was reversed. To quantify the apparent preferential binding, we varied the tau concentration and measured the resulting binding to the separate regions of the microtubule (Figure 1C). We found that tau’s affinity for GDP-tubulin extensions were ∼5-fold higher than for GMPCPP seeds. As shown in ^7^, we found that tau had a slight preference for GTPγS microtubules over GMPCPP microtubules, although the affinity was still lower than that of GDP microtubules (Figure S1). Therefore, we conclude that tau has the ability to recognize different tubulin nucleotide states and binds to GDP-tubulin in preference to GMPCPP-tubulin in microtubules.

**Figure 1.**
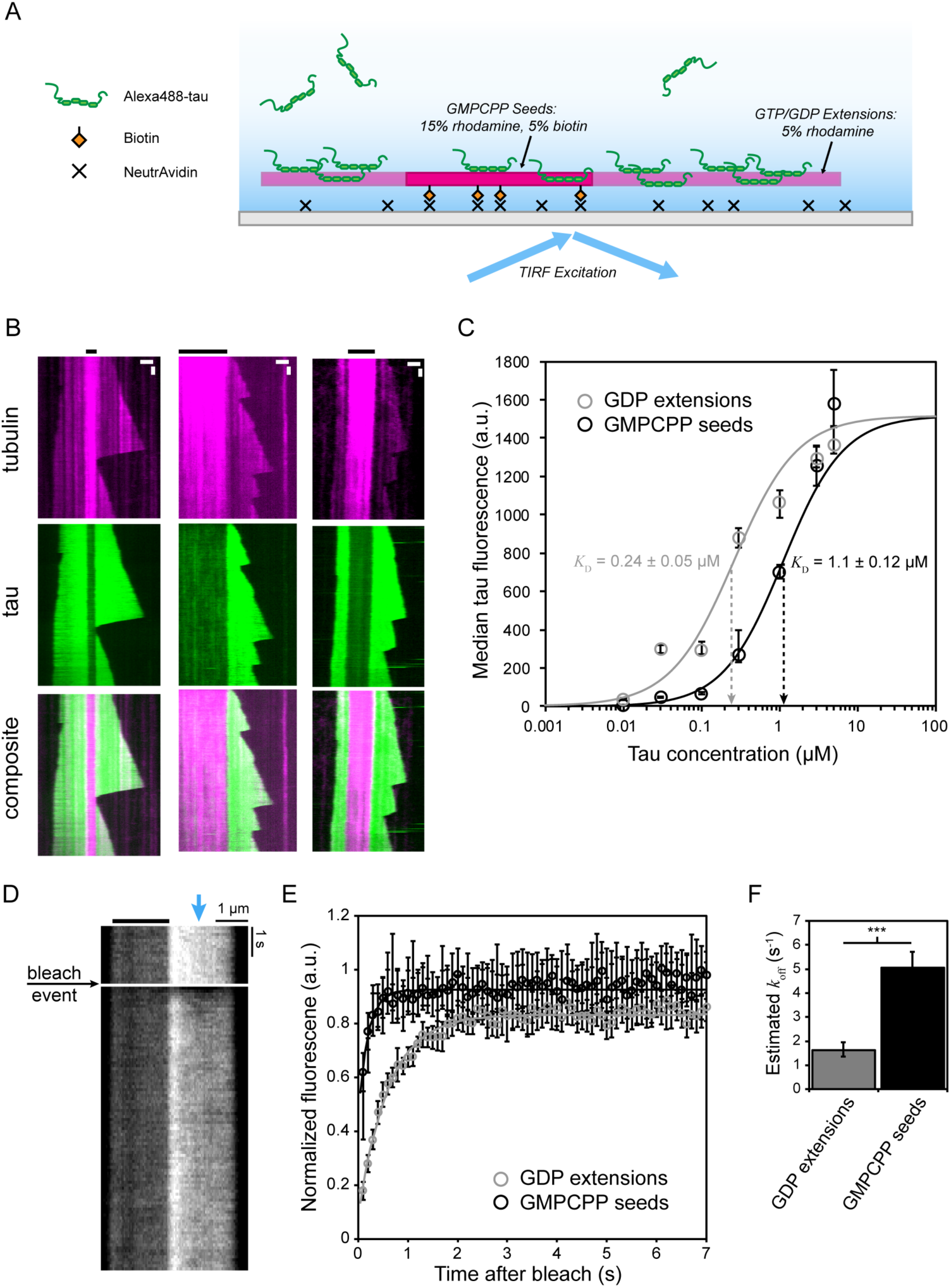
2N4R tau preferentially binds microtubules composed of GDP-tubulin over GMPCPP-microtubules *in vitro*. A) Diagram of the TIRF microscopy assay used to measure tau binding to microtubules. Bright (15% rhodamine, 5% biotin labeled) GMPCPP seeds were conjugated to the coverslip using NeutrAvidin. Dim (5% rhodamine label) GTP/GDP extensions were grown from seeds in the presence of Alexa488-tau. B) Kymographs of individual microtubules grown in the presence of 300 nM tau. Black bars indicate the location of the GMPCPP seeds. Scale bars are 1 µm and 30 s in the x- and y-direction, respectively. C) Binding of Alexa488-tau to microtubule seeds (black) and extensions (gray) under a range of tau concentrations. Curves indicate best-fit Hill function to all data points. Value of *K*_D_ ± 95% confidence interval (CI) resulting from the fit is shown. Error bars are ± 95% CI. D) Kymograph showing the fluorescence recovery after photobleaching (FRAP) of Alexa488 labeled tau bound to the GDP extension. Blue arrow is the center of the focused laser spot. Thick black bar indicates the location of the GMPCPP seed. E) FRAP recovery curves for seeds (black) and extensions (gray) in the presence of 300 nM tau. Lines are best fit to Eq. 2. F) Mean off-rate constant estimated from fits to the FRAP recovery curves for each individual microtubule. Data from both 300 nM and 1 µM tau were combined in estimating the mean for GMPCPP seeds. Error bars are ± 95% CI.

The observed higher GDP-tubulin binding affinity could be a result of a faster on-rate and/or slower off-rate from GDP-tubulin. To further explore tau’s nucleotide preference we photobleached Alexa488-tau on different regions of the microtubule and measured the fluorescence recovery after photobleaching (FRAP) (Figure 1D). As seen in Figure 1E, fluorescence recovery occurred faster on GMPCPP seeds as compared to GDP-tubulin extensions. Under the experimental conditions, recovery is determined by the off-rate constant^12^, and thus tau has a higher off-rate from GMPCPP-tubulin than for GDP-tubulin. Furthermore, the fold difference is approximately equal to the observed fold shift in affinity. Therefore, we conclude that tau’s increased affinity for GDP-tubulin is a result of a reduced unbinding rate constant.

### Tau is less abundant at the growing end of microtubules *in vitro*

Since GMPCPP is an analog of GTP, it is possible that tau merely prefers GMPCPP and binds to microtubule-bound GTP-tubulin with similar or even greater affinity to that of tau binding to GDP-tubulin. If tau indeed prefers GDP-over GTP-tubulin, then it should be absent from the tip of growing microtubules, the exact opposite of the characteristic EB enrichment on the tips of growing microtubules. To test if this was the case, we tracked the end of dynamic microtubules using our previously published tracking procedure^13, 14^ and quantified the average fluorescence along the microtubule axis. To improve tracking precision, the rhodamine-tubulin ratio was reversed such that 15% of the tubulin in solution was labeled (Figure 2A). This labeling reversal did not influence the observation of tau enrichment on GDP-extensions versus GMPCPP seeds (Figure 2B). As seen in Figure 2C, the leading edge of tau fluorescence, on average, lagged behind the leading edge of tubulin fluorescence leading to a statistically significant (*p* < 0.001) offset of 78 nm ± 15 nm (median ± 95% CI) between the two during periods of growth. This observation is consistent with tau having lower affinity to the stabilizing cap of GTP-tubulin as compared to regions of GDP-tubulin proximal to the microtubule tip. The offset is also consistent with previous estimates of the stabilizing cap size *in vitro*^5, 15^.

**Figure 2.**
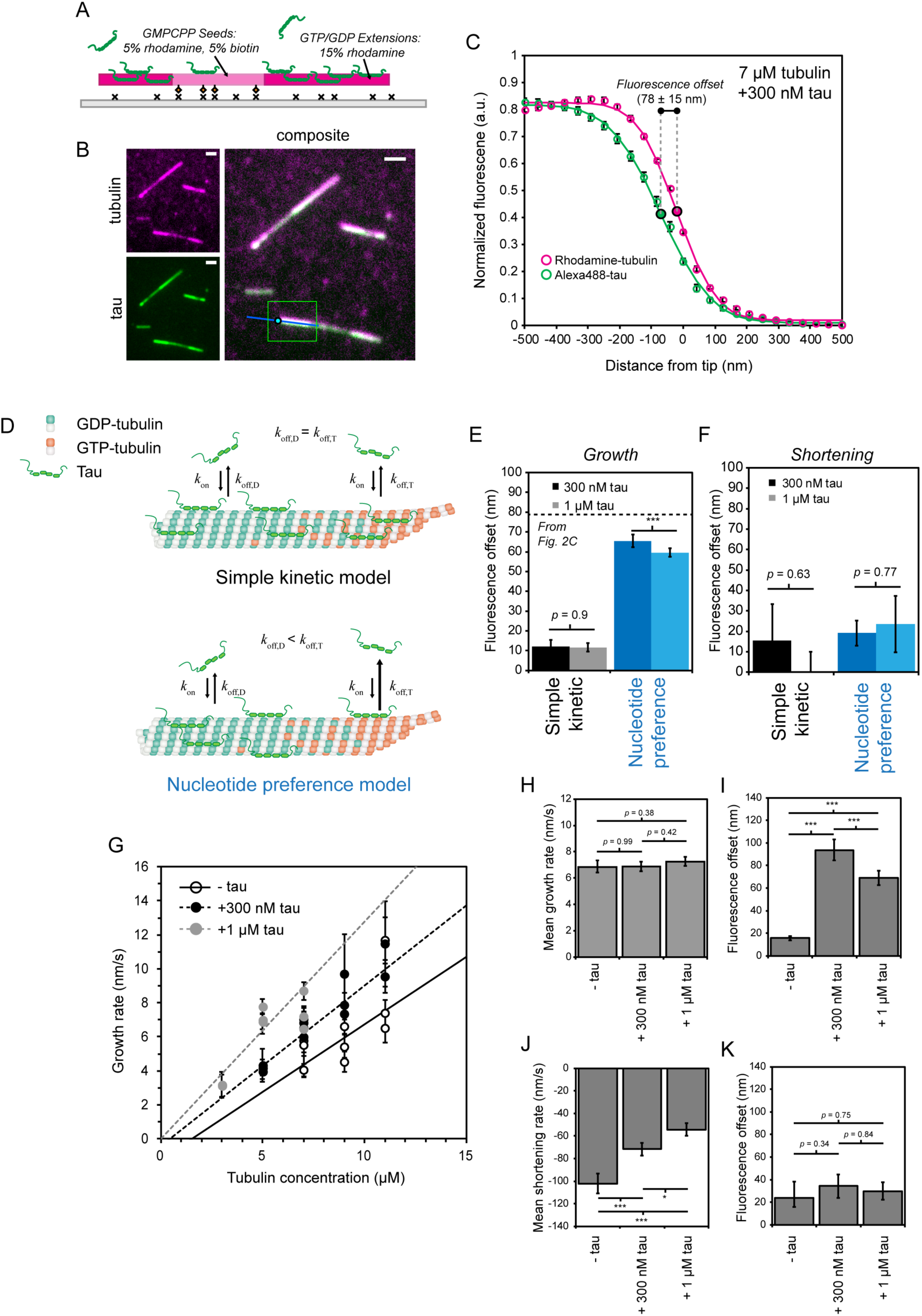
Tau avoids the growing microtubule GTP cap *in vitro*. A) TIRF microscopy diagram showing the rhodamine labeling ratios of the microtubule seeds and extensions used for tip tracking. To improve tip tracking precision, rhodamine tubulin labeling percentages for seed and extensions were switched. B) Example microtubules imaged by TIRF microscopy and used for tip tracking and fluorescence profile analysis. Green box indicates the region of interest used in TipTracker. Blue line is the determined microtubule axis and cyan dot is the estimated microtubule tip position. C) Average fluorescence along the microtubule axis during periods of growth in the presence of 7 µM tubulin and 300 nM Alexa488-tau. Fluorescence values were centered using the tip position estimated from the rhodamine-tubulin signal using TipTracker. Lines are best fit of a Gaussian survival function, Eq. 1. Error bars are ± SEM. Fluorescence offset value indicated is the median ± 95% CI. D) Simulated tau binding models. In a simple kinetic model (top), tau binds and unbinds from all sites independent of nucleotide state. In the nucleotide preference model, tau’s off-rate constant is higher for GTP-tubulin (orange) compared to GDP-tubulin (green). E-F) Simulation results for each model in the presence of 300 nM (dark) and 1 µM (light) tau, simple kinetic model is in black while the nucleotide preference model is shown in blue. Dashed line (E) is the median fluorescence offset value from Figure 2C above for 7 µM tubulin + 300nM tau. Error bars are SEM. G) Average growth rate as a function of tubulin concentration under three conditions, without tau (open circles), +300 nM tau (closed black circles), and +1 µM tau (closed gray circles). Individual dots are the means for individual assay preparations. Error bars are ± 95% CI. Lines are linear best fit to all microtubules. H) Mean microtubule growth rate for all microtubules analyzed, independent of tubulin concentration. Error bars are ± 95% CI. I) Mean offset between rhodamine-tubulin and Alexa488-tau fluorescence signals during growth, as determined by individual fits of Eq. 1. Error bars are ± 95% CI. J) Mean microtubule shortening rate for all microtubules analyzed, independent of tubulin concentration. Error bars are ± 95% CI. K) Mean offset between rhodamine-tubulin and Alexa488-tau signals during shortening. Error bars are ± 95% CI. * *p* < 0.05, ** *p* < 0.01, *** *p* < 0.001.

As the microtubule grows, and tubulin subunits are added to the microtubule lattice, it will necessarily take time for tau to bind to newly created binding sites. If the rate of tau on-off kinetics are slow compared to site creation, then tau binding will lag behind the growing end. Therefore, it is feasible that the observed offset is simply a result of tau on-off kinetics, as opposed to a binding preference for GDP-tubulin. To test which model is more consistent with experimental observations, we simulated tau’s association dynamics with the microtubule under the two different models (Figure 2D). In the simple kinetic model (Figure 2D top), tau is allowed to bind to any site on the microtubule with uniform affinity. Here, tau’s simulated off-rate and binding affinity was equal to that measured experimentally for microtubule extensions (Figure 1). In the second model, nucleotide preferential binding (Figure 2D bottom), tau’s on-rate was uniform for all sites on the microtubule but the off-rate for GTP-tubulin was based on the faster rate observed for GMPCPP seeds experimentally (Figure 1E-F). To compare simulation output to experimental measurements, we used model convolution^16^ to simulate resulting tubulin and tau images, which were then processed through the same tip tracking algorithm in order to estimate any potential offset between the tau and tubulin fluorescence. We found that the simple kinetic model predicted a smaller offset compared to the nucleotide preference model during periods of growth that was quantitatively smaller than the offset observed experimentally (Figure 2E). In the nucleotide preference model, increasing the tau concentration from 300 nM to 1 µM, closer to the *K*_D_ of GMPCPP-tubulin (or GTP-tubulin in the model), resulted in a slightly decreased offset (Figure 2E). This was a result of tau’s increased binding to regions of GTP-tubulin at the microtubule tip. In the simple kinetic model, the offset was not influenced by the tau concentration, although this simply reflects the low offset initially at 300 nM. The observed offset in the nucleotide preference model was predicted to be eliminated during periods of shortening (Figure 2F). Thus, the model predicts that if tau indeed binds to GTP-tubulin with similar affinity to that of the analog GMPCPP-tubulin, then an experimental offset during growth should be of similar magnitude and be reduced by an increase in tau concentration. Furthermore, any potential offset should be lost during shortening.

We tested the model predictions in our *in vitro* assay in the presence of 300 nM and 1 µM Alexa488-tau. As seen in Figure 2G, tau moderately increased the growth rate *in vitro*, similar to previous observations^8^. Because EB comet size (or GTP cap size) is proportional to the rate of growth^17^, this adds a potential complication to the analysis at a single tubulin concentration. For example, if microtubules grow faster in the presence of 1 µM tau compared to 300 nM or no tau, then the GTP cap size will be larger, thereby increasing the potential offset in a nucleotide preference model. Therefore, we compensated for this effect on growth rate by adjusting the tubulin concentration in the presence of tau such that the average microtubule growth rate was equal across all conditions (Figure 2H). Experimentally, we found that the offset in the presence of 300 nM tau was comparable to that predicted from the nucleotide preference model, but quantitatively inconsistent with the simple kinetic model (Figure 2I). Additionally, as predicted by the nucleotide preference model, the experimental offset decreased in the presence of 1 µM tau compared to 300 nM. The experimental shortening rate was reduced with increasing tau concentrations (Figure 2J), suggestive of a stabilizing effect of tau. As predicted by the nucleotide model, the observed fluorescence offset during growth was lost during periods of shortening (Figure 2K). Thus, experimental offset trends were qualitatively and quantitatively consistent with the nucleotide preference model and inconsistent with the simple kinetic model. In summary, the simple kinetic model is incapable of producing an offset comparable to that observed experimentally. Therefore, we conclude that tau is offset from the microtubule plus-end due to preferentially binding GDP-tubulin over GTP-tubulin *in vitro*.

### Tau is a microtubule tip avoider in mammalian cells

To determine if tau exhibits a nucleotide preference in living cells we transiently expressed human 2N4R tau tagged with EGFP along with mCherry-α-tubulin in LLC-PK1 mammalian cells. While these cells are not neuronal, they provide multiple advantages over neurons. First, they form expansive, thin regions where single microtubules are readily visible and are ideal for nanoscale quantitative analysis^18^. Additionally, since epithelial cells do not natively express tau, all tau in the cells should be tagged with EGFP. We found that tau-EGFP was expressed at variable levels and that it was localized to microtubules (Figure 3A). Additionally, tau-EGFP exhibited binding characteristics consistent with those previously observed^11^, in particular tau was enriched on curved regions of the microtubule at low expression levels (Figure 3A, arrows).

**Figure 3.**
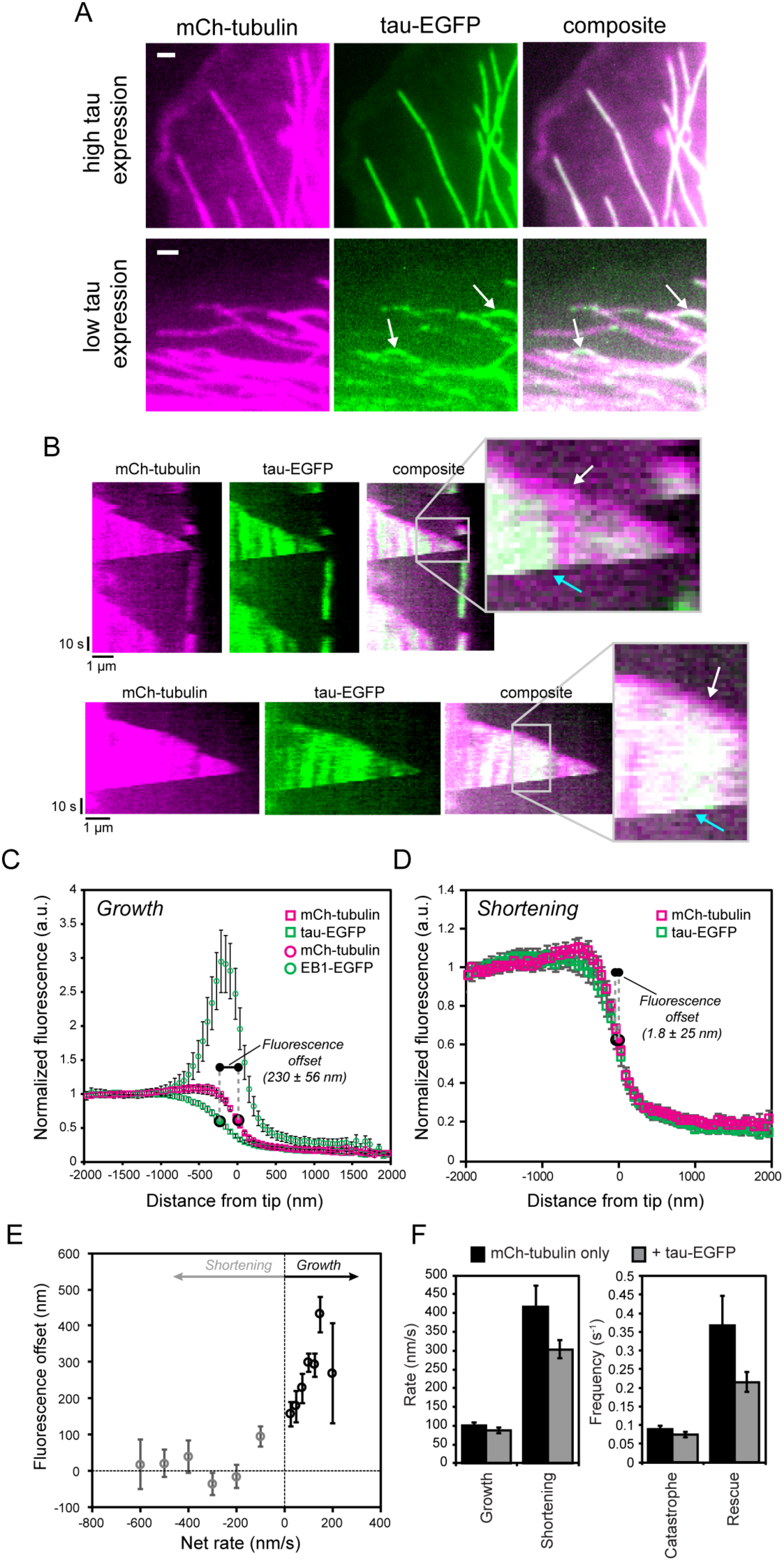
Tau avoids the growing microtubule GTP cap *in vivo*. A) LLC-PK1 cells expressing mCherry-tubulin and 2N4R tau-EGFP with varying tau-EGFP expression levels. Arrows indicate tau enrichment on curved regions of microtubules at low expression levels. Scale bars are 1 µm. B) Example kymographs showing microtubule dynamics in the presence of tau-EGFP. White arrows indicate periods of growth while cyan arrows indicate periods of shortening. C-D) Average fluorescence signal of mCh-tubulin, tau-EGFP, and EB1-EGFP along the microtubule axis during periods of growth (C) or shortening (D). Fluorescence values were centered using the tip position estimated from the mCherry-tubulin signal using TipTracker. Alike symbols (squares or circles) indicate paired fluorophores for dual expression. Fluorescence offset values indicated are the median ± 95% CI. Error bars are ± SEM. E) Fluorescence offset as a function of microtubule net assembly rate. Values were binned based on the net-rate. Offset values for growth periods are in black and those for shortening periods are in gray. Error bars are ± 95% CI. F-G) Dynamic instability parameters in cells expressing mCherry-tubulin only (black) or mCherry-tubulin and tau-EGFP (gray). Error bars are ± 95% CI.

+TIPs are characterized by several features in living cells. First, they are enriched at the microtubule plus end during growth and absent during shortening, and, second, the amount of protein at the microtubule plus end is positively correlated with the rate of microtubule growth. We expect that if tau is acting as a microtubule tip avoider, then it should exhibit the opposite characteristics; it will be offset from the microtubule plus end during growth but not shortening, and this offset should be positively correlated with the rate of microtubule growth. Consistent with a tip avoider, we observed a separation of the tubulin signal from that of tau-EGFP during periods of growth in cells co-expressing tau-EGFP and mCherry-α-tubulin (Figure 3B, white arrow). This visible offset subsequently disappeared during periods of shortening (Figure 3B, cyan arrow). To further quantify this observation, we performed the same tau and tubulin fluorescence profile analysis used *in vitro* in the LLC-PK1 cells, and compared it to the fluorescence profile of EB1. As seen in Figure 3C, tau was, on average, offset from the plus end of the microtubule *in vivo* during periods of growth. Interestingly, the initial drop in tau-EGFP fluorescence was directly aligned with the initial rise in EB1-EGFP fluorescence (Figure 3C). Additionally, we found that the average offset (226 ± 38 nm; mean ± 95% CI) was consistent with our previous estimates of the GTP cap size in the same cells, 750 GTP-tubulin subunits or 310 nm^5^. During periods of shortening, the tau and tubulin fluorescence converged such that there was no longer an offset (Figure 3D), as observed in the kymographs. *In vivo*, microtubules exhibit a much wider range of growth rates compared to *in vitro*. To further test our hypothesis, we binned microtubules by growth rate and plotted their rate versus average fluorescence offset. Consistent with the prediction for a tip avoider, fluorescence offset was directly correlated with growth rate, increasing for faster growing microtubules (Figure 3E). Furthermore, an offset during shortening was not detected and was not correlated with rate of shortening. As a result of being absent from the microtubule plus end during growth, we did not observe any effect of tau-EGFP expression on microtubule growth rate or catastrophe frequency (Figure 3F). While we did not observe an offset trend with respect to growth *in vitro* (Figure S2A), this was likely a result of the narrow range of growth rates observed *in vitro* compared to *in vivo*. The overall trend of net rate versus offset *in vitro* was comparable to that observed *in vivo* (Figure S2B). Based on these results, we conclude that tau acts as a microtubule tip avoider, absent from the microtubule plus-end, in mammalian cells.

### Tau offset *in vivo* is consistent with a preferential binding model

Unlike *in vitro*, it is possible that other MAPs in LLC-PK1 cells interfere or compete with tau for binding sites, resulting in the observed offset in tau fluorescence from the microtubule plus end. EB1, in particular, could be such a competitor as it binds preferentially to the microtubule plus end during growth and is expressed at high concentrations^5^. To test whether a competitive binding model could be consistent with experimental observations *in vivo*, we first quantified the tau-EGFP binding kinetics. FRAP of tau bound to regions of the microtubule resulted in an estimated off-rate constant of 2.02 s^-1^ (Figure 4A), consistent with that estimated for GDP microtubules *in vitro*. To construct a tau binding curve, we estimated the amount of tau-EGFP bound per length of microtubule using our previous approach for quantifying EB1 concentrations in the same cells^5^. All regions analyzed were proximal to the microtubule plus end (i.e. towards the minus end) and presumably consisted of GDP-tubulin. We found that tau-EGFP bound to microtubules with *K*_D_= 1.1 µM, which was qualitatively higher than our estimate for GDP-tubulin *in vitro*, although not statistically significant due to the uncertainty in the *in vivo* measurement. From these estimates we constructed a computational model for *in vivo* tau binding kinetics similar to the *in vitro* model (Figure 2C-D). In addition to the simple kinetic model and nucleotide preference model, we constructed a third model where EB1 and tau compete for binding sites (Figure 4D). Values used for EB1 kinetics and concentration were from our previous estimates in these cells^5^. We found that the only model consistent with *in vivo* experimental observations of tau binding was the nucleotide preference model (Figure 4F-G).

**Figure 4.**
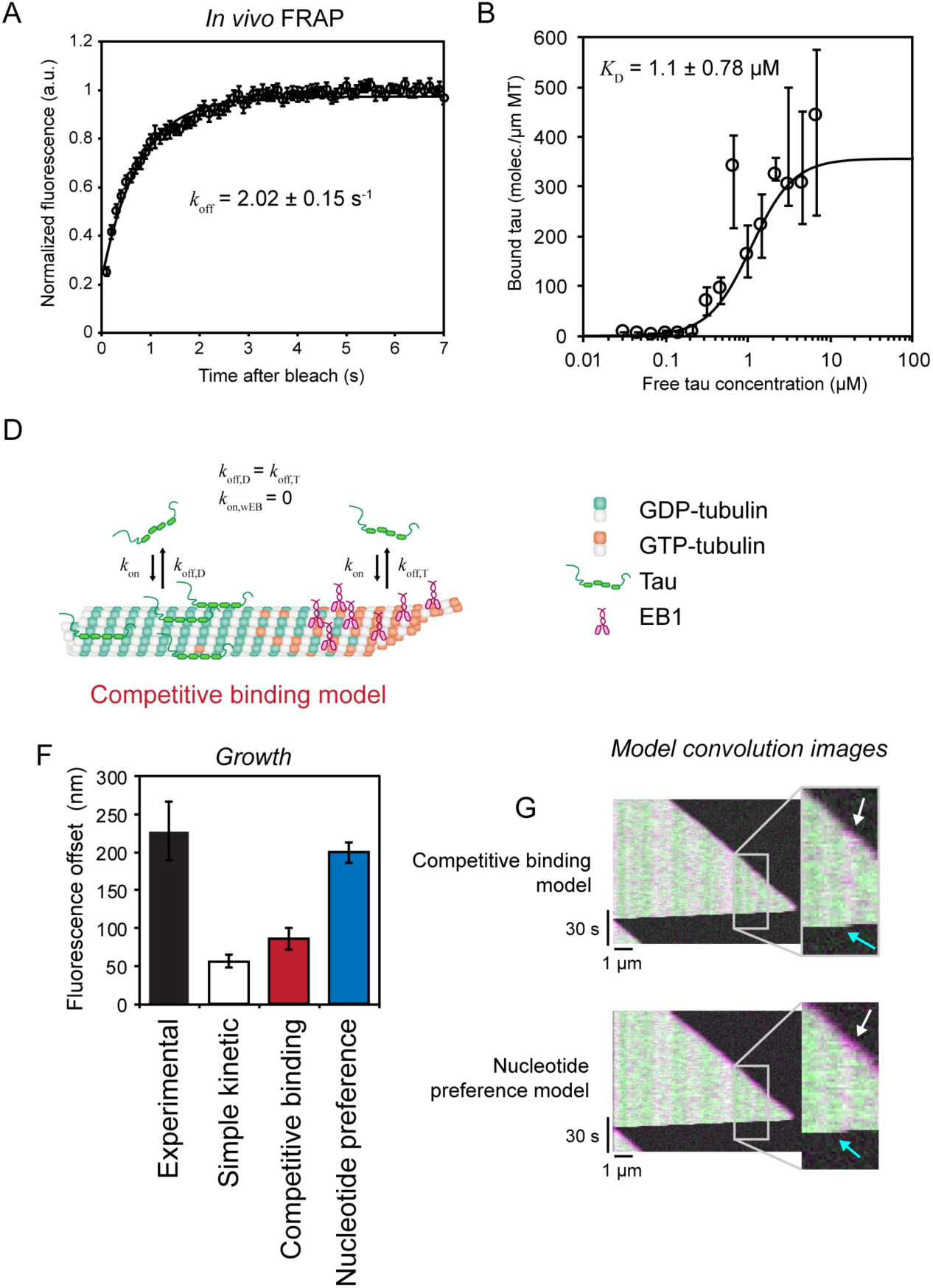
Tau offset from the growing plus end cannot be explained by tau-GFP dynamics or a competitive binding model, but is consistent with intrinsic nucleotide preference for GDP-tubulin over GTP-tubulin. A) FRAP of tau-EGFP bound to microtubules, proximal to the microtubule tip, in cells expressing mCherry-tubulin and tau-EGFP. Line is best fit exponential recovery. Error bars are ± 95% CI. B) Tau bound per length of microtubule versus the free tau-EGFP concentration estimated from the background intensity levels. Line is best fit Hill function to all data points. Data was binned based on free tau concentration values. Error bars are ± 95% CI. C-E) Diagrams for different tau binding models used in simulation. C) A simple, non-competitive binding model where tau binds with equal affinity to all sites on the microtubule. D) A competitive binding model in which tau competes with EB1 for sites along the length of microtubule. Tau is not allowed to bind to sites occupied by EB1 (*k*_on,wEB_ = 0). EB1 binds with higher affinity to GTP-tubulin compared to GDP-tubulin, which gives it microtubule plus end tracking ability. E) A nucleotide preference model in which tau binds with higher affinity to GDP-tubulin compared to GTP-tubulin. As in the competitive model, tau is not allowed to bind to sites occupied by EB1. F) Model-predicted offset as a function of the simulated tau kinetics. Error bars are ± 95% CI. G) Simulated images of tubulin (red) and tau (green) fluorescence signal resulting from the models in C and E. Microtubule dynamics were simulated using the *in vivo* parameters from^18^. White and cyan arrows identify periods of growth and shortening, respectively.

Neither the simple model nor the competitive binding model were able to predict experimental observations of the tau fluorescence offset (Figure 4F). Additionally, simulated images obtained through model convolution of the model output^16^ showed that the nucleotide preference model resulted in microtubule tip avoider behavior, where a distinguishable separation between tau and tubulin fluorescence was detectable during growth (Figure 4G, white arrows) and subsequently absent during shortening (Figure 4G, cyan arrows), consistent with experimental observations in the presence of tau (Figure 3B). This visible offset was not seen in either the simple model or the competitive binding model. Based on these observations, we conclude that tau acts as a microtubule tip avoider *in vivo* by preferentially binding GDP-tubulin over GTP-tubulin.

## Discussion

By performing high resolution, quantitative fluorescence measurements of tau binding to microtubules *in vitro* and *in vivo* we show here that tau avoids the GTP cap at the growing end of microtubules, resulting in an offset or lag of tau at the growing plus end. Additionally, the size of this offset is positively correlated with the microtubule growth rate, the exact opposite of +TIP tracking proteins. We find that these observations are consistent with a nucleotide preference model, in which tau preferentially binds unstable GDP-tubulin over the more stable form of GTP-tubulin. To our knowledge, the evidence that we show here is the first example of a MAP that preferentially binds GDP-tubulin, thereby functioning as a growing microtubule tip avoiding protein.

One interesting potential benefit to being a tip avoiding MAP is that tau could influence microtubule dynamics specifically during shortening, thus perhaps controlling the extent of microtubule loss during shortening without interfering with the dynamics of growth and catastrophe. This would allow microtubule plus ends to continue to dynamically explore the intracellular space while controlling the extent of large-scale losses of polymer mass that can occur during the shortening phase. It is interesting to speculate that this capability would allow neuronal microtubule plus ends to dynamically grow into and shorten from axonal branches and dendritic spines without risking large scale depolymerization-mediated microtubule polymer loss from axons and dendrites.

Tau has long been believed to act as a microtubule stabilizer and promote microtubule bundling in axons, although the stabilizing role of tau has recently been challenged^19^. The aberrant regulation of tau phosphorylation states causes it to dissociate from microtubules and self-associate, forming neurofibrillary tangles characteristic of tauopathies such as Alzheimer’s disease and dementia. It is possible that tau hyperphosphyorylation and/or mutation in these neurological diseases exert their phenotype by their failure to properly recognize GDP-tubulin. The vast majority of tubulin in microtubules is GDP-bound, and such a loss of function would significantly shift the distribution of tau in neurons. Furthermore, more tau would be available to bind to GTP-tubulin, thus displacing +TIPs, such as EB3, and their associated activities into abnormal locations in the neuron.

One remaining question, however, is what is the specific mechanism by which tau distinguishes the tubulin nucleotide states? Recently, cryo-EM has shown the structural differences of GDP- and GTP-tubulin in the microtubule lattice, specifically that GTP-tubulin undergoes a compaction at the longitudinal interface upon nucleotide hydrolysis^20^. This compaction likely leads to a change in the intradimer distance between tubulin subunits, where tau binds to the microtubule^21^, that is energetically favorable to tau binding. This could also explain observations that tau preferentially binds curved regions of microtubules^11^, where the intradimer distance will be reduced on the inside of the microtubule curves compared to straight regions. Furthermore, preliminary data suggests that tau’s nucleotide binding preference is specific to tubulin in microtubules and is not present when binding to tubulin in solution (Figure S3). Thus, it appears that a possible explanation for tau’s nucleotide preference is that it is reading out the structural changes in the microtubule lattice structure that are a result of hydrolysis.

## Materials and Methods

### Tau cloning, purification, and labeling

The parent tau plasmid encodes for the longest tau isoform, 2N4R tau. It includes an N-terminal His-tag with a tobacco etch virus (TEV) protease cleavage site for purification^22^. The native cysteines, C291 and C322, are mutated to serine to allow for the introduction of cysteine at residue S433 for site-specific labeling. This mutation does not impact microtubule polymerization *in vitro*^23^.

Tau protein expression was induced with 1mM isopropyl β-D-1-thiogalactopyranoside (IPTG) at OD ∼0.6 overnight at 16 °C. Purification was based on previously reported methods^22^. Briefly, cells were lysed by sonication, and the cell debris pelleted by centrifugation. The supernatant was incubated with Ni-NTA resin (Qiagen or BioRad) and the recombinant protein was bump eluted with 500 mM imidazole. The His-tag was removed by incubation with lab purified TEV proteinase overnight at 4 °C (constructs >200 residues). Uncleaved protein was removed by a second pass over the Ni-NTA column. Remaining contaminants were removed using size exclusion chromatography on a HiLoad 16/600 Superdex 200 Column (GE Life Sciences, Marlborough, MA). Proteins that did not require fluorescent labeling were buffer exchanged using Amicon concentrators (MilliporeSigma, Burlington, MA) into storage buffer (20 mM Tris pH 7.4 and 50 mM NaCl) and snap frozen for storage at −80°C.

Site specific labeling of tau for fluorescent measurements was carried out as described previously^22^. Briefly, tau was incubated with 1 mM dithiolthreitol (DTT) for 30 minutes, and then buffer exchanged into labeling buffer (20 mM Tris pH 7.4, 50 mM NaCl, and 6 M guanidine HCl). Alexa Fluor 488 maleimide was added in 2-fold molar excess and incubated at room temperature for 10 minutes, followed by overnight incubation at 4°C. Labeling reactions were protected from ambient light and with constant stirring; the dye was added dropwise. The labeled protein was buffer exchanged into 20 mM Tris pH 7.4 and 50 mM NaCl and unreacted dye was removed using HiTrap Desalting Columns (GE Life Sciences). Labeled protein was then snap frozen for storage at −80°C. The addition of Alexa488 did not influence tau-induced tubulin polymerization overall (Figure S4).

### Preparation and functionalization of imaging chambers

Imaging chambers for TIRF microscopy were assembled and functionalized as described in^24^, with few modifications. First, an acid cleaned coverslip^25^ was rendered hydrophobic by brief incubation in Rain-X® Original Glass Water Repellent (ITW Global Brands, Houston TX) at room temperature. Coverslips were then allowed to dry completely before remaining residue was wiped away using lens paper. Hydrophobic coverslips were then mounted to acid cleaned glass slides using double sided tape, forming three separate imaging channels.

Imaging chambers were functionalized by flowing in solutions of 0.1 mg/mL NeutrAvidin (ThermoFischer Scientific, Waltham, MA) in PBS, followed by 5% Pluronic® F-127 (Sigma-Aldrich, St. Louis, MO) in PBS, and then doubly stabilized GMPCPP microtubule seeds (5% or 15% rhodamine-labeled, 5% biotin-labeled) in BRB80 (80 mM PIPES/KOH pH 6.9, 1 mM EGTA, 1 mM MgCl_2_). For each separate solution, chambers were allowed to incubate at room temperature for 10 mins and then washed with 8-10x chamber volume of BRB80. After the last wash, 2x chamber volumes of the imaging solution was flowed through the imaging chamber before moving to the microscope for imaging. Imaging solution consisted of indicated concentration of porcine brain tubulin (5% or 15% rhodamine-labeled, Cytoskeleton Inc., Denver CO) and Alexa488-labeled tau in BRB80 supplemented with 1 mM GTP, 40 mM D-glucose, 8 µg/mL catalase, 20 µg/mL glucose-oxidase, and 0.1 mg/mL casein. To prevent sample drying during imaging, individual imaging chambers were sealed using CoverGrip sealant (Biotium, Fremont, CA).

### TIRF microscopy

Unless otherwise noted, all proteins were handled and stored as described previously^24^. Rhodamine-labeled microtubules growing from double-stabilized GMPCPP microtubule seeds were imaged by TIRF microscopy using a 100x, 1.49NA Apo TIRF objective on a Nikon TiE inverted stand equipped with the Perfect Focus, H-TIRF module and LU-N3 laser launch (Nikon Instruments Inc., Melville, NY) under control of NIS-Elements software (v4.xx, Nikon Instruments). Images were collected on a Zyla 4.2 PLUS sCMOS camera (Andor, Belfast, UK) with a high speed emission filter wheel (HS-632; Finger Lakes Instrumentation, Lima, NY) placed between the camera and stand for color separation. Additional 1.5x tube lens in the microscope stand resulted in a total magnification of 150x (42 nm/pixel). For tau binding and microtubule dynamics imaging, 488 nm and 561 nm TIRF lasers were reflected up through the rear aperture of the objective using a triple band pass filter set (TRF69901; Chroma Technology Corp., Bellows Falls, VT). Unless otherwise noted, all images were collected using 200 ms exposure at 20% laser power. For FRAP experiments, a 488 nm 100 mW Argon-ion laser (Spectra-Physics, Santa Clara, CA) shuttered by a Uniblitz VS35 shutter (Vincent Associates, Rochester, NY) was focused on the imaging plane as previously described^18^ using a separate light path from that used for TIRF imaging. Bleach event timing was set to a 3 s delay and 100 ms exposure using a VMM-TI shutter driver/timer (Vincent Associates). Simultaneous TIRF imaging was accomplished by replacing the triple band pass filter above with an 80/20 beam splitter. To compensate for the beam splitter, the 488 nm TIRF laser was increased to 100% power such that 20% laser power used for imaging was maintained. Temperature was maintained at 37°C using an objective heater (OkoLab S.R.L., Pozzuoli, Italy) and airstream incubator (Nevtek, Burnsville, VA).

### Live cell imaging

LLC-PK1 cells used for imaging were cultured in glass-bottomed 35 mm dishes as previously described^18^. After 24 hrs in culture, LLC-PK1 cells were transiently transfected with 2N4R tau-EGFP and mCherry-α-tubulin using FuGENE® HD transfection reagent (Promega, Madison, WI) according to the manufacturer’s instructions. Briefly, cells were incubated overnight with a 4:1 FuGENE:pDNA ratio in OptiMEM+10% FBS. The same FuGENE:pDNA ratio was maintained for cells transiently transfected with mCherry-α-tubulin only. Prior to imaging, FuGENE and pDNA were removed from the sample by media exchange.

Transiently transfected LLC-PK1 cells were imaged on the same Nikon TiE stand (Nikon Instruments) used for TIRF. A 100x 1.49NA Apo TIRF objective (Nikon Instruments) without additional tube lens resulted in a final pixel sampling size of 65 nm. Images were collected at 500 ms intervals for a total of 1 min using 200 ms exposure under illumination from SpectraX Light Engine (Lumencor, Beaverton, OR) at 50% power. The stage and objective were maintained at 37°C for the duration of imaging using environmental control provided by a BoldLine stagetop incubation system (OkoLab).

### Tau binding simulations

*In vitro* and *in vivo* microtubule assembly dynamics were simulated as previously described^18^, using the pseudomechanical model^26^. Since the tau binding profile was the simulation output of interest, microtubule dynamics simulations were run prior to and independent of tau binding simulations. Tau on-off kinetics were simulated for all subunits in the microtubule lattice based the probability of tau binding or unbinding as determined by *p* = 1-exp(-*k*\**t*), where *k* is tau’s on-rate (*k*\*_on_ = *k*_on_*[*tau*]) or off-rate (*k*_off_ = *k*_on_\**K*_D_; *K*_D_ is tau’s affinity for the microtubule) and *t* is the current time step from the microtubule simulation. After determining the on-off probabilities, a random number, *r_i_*, was generated for each individual subunit and the event was executed if *r_i_* < *p*. The execution order for binding and unbinding was randomly determined at each time step, *t.* For the simple kinetic model, it was assumed that tau kinetics were equivalent for all sites on the microtubule. For the nucleotide preference model, tau’s off-rate was determined by the nucleotide state of the tubulin subunit that it was associated with; here *k*_off,T_ = *k*_on_\**K*_D,T_, *k*_off,D_ = *k*_on_\**K*_D,D_, and *K*_D,D_ < *K*_D,T_. EB1 binding kinetics were handled in the same manner as tau using concentration and *K*_D_ values estimated previously^5^. For the competitive binding model, tau and EB1 binding was not permitted to sites already occupied by either protein. To quantify fluorescence offset, simulation output was model convolved^16^ based on the experimental imaging setup to create simulated fluorescence images. These model convolution images were then run through the same analysis scheme used for experimental images. All simulations were carried out in MATLAB R2017a (The Mathworks Inc, Natick, MA).

### Microtubule tip tracking and fluorescence profile analysis

The dynamic microtubule end was tracked using our previously described semi-automated algorithm^13, 14^, TipTracker (version 3.1), without modification. Briefly, fluorescence profiles along the determined microtubule axis (x’’-axis) are fit with a Gaussian survival function

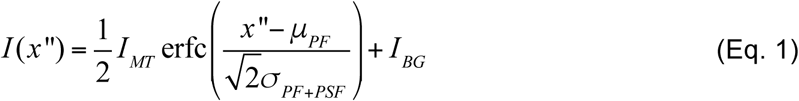

where *µ*_PF_ is taken to be the position of the microtubule tip, *I*_MT_ and *I*_BG_ are the fluorescence intensity on the microtubule and the background, respectively, and *σ*_PF+PSF_ is the spread of the fluorescence due to the combination of the point spread function and the taper or spread of protofilament lengths at the microtubule tip. For averaging purposes, fluorescence profiles along the microtubule axis from all channels (tubulin, tau, and EB1) were aligned to *µ*_PF_ determined from the tubulin channel (either rhodamine-tubulin or mCherry-α-tubulin). Fluorescence offset values were the difference between *µ*_PF_ resulting from tracking the tubulin and tau channels (offset = *µ*_PF,tub_ – *µ*_PF,tau_). *In vitro* fluorescence profiles were normalized to the maximum value while *in vivo* profiles were normalized to an average of the first 10 values.

For the current imaging setup, the fluorescence intensity per EGFP molecule per exposure was determined using LLC-PK1α cells stably expressing EGFP-α-tubulin^27^ as previously described^5^. This value was then used to estimate the number of tau molecules bound per length of microtubule by dividing the integrated fluorescence by the length of the microtubule segment. Since LLC-PK1 cells are epithelial and do not express tau, it was assumed that all tau molecules in the cells expressing tau-EGFP were tagged.

FRAP curves were fit with an exponential recovery curve by weighting early time points during recovery more strongly than steady-state values as described previously^18^. Prior to fitting, FRAP values were corrected using microtubules outside the bleached region and then normalized to the average value over 2 s (20 frames) prior to the bleach event. Assuming the tau dynamics are at steady-state with microtubules *in vitro* and *in vivo*, and that the amount of bleached tau is small compared to the total amount of tau, then the exponential recovery is dictated by the off-rate as described in^12^;

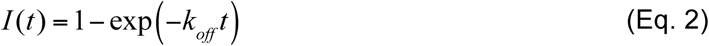

where *k*_off_ is the tau unbinding rate.

### Statistical analysis

Unless otherwise noted, comparisons between experimental conditions were performed by one-way analysis of variance (ANOVA) statistical tests performed in MATLAB (R2017a) using the *anova1* function, and where appropriate, corrected for multiple comparisons using the *multcompare* function. For all statistics, 95% confidence intervals were estimated using a bootstrapping method as previously described ^18^.

## Acknowledgements

Research reported in this publication was supported by the National Institute of Aging of the National Institutes of Health under award number RF1AG053951 to E.R and D.J.O. and Institutional Training Grant T32 GM071399 to K.M.M. The authors thank Virginia Lee at the University of Pennsylvania and Lynne Cassimeris at Lehigh University for providing fluorescent protein plasmids and also thank Jonathan Sachs and Mahya Hemmat for helpful discussions.

## Code availability statement

All code used for simulations and analysis is available upon reasonable request to the corresponding author.

## Author contributions

All experiments and analysis were performed by B.T.C. B.T.C. and D.J.O. conceived and designed the experiments. K.M.M. purified and labeled full-length tau protein. E.R. and D.J.O. supervised the study. B.T.C wrote and D.J.O edited the manuscript. All authors contributed to data interpretation, and commented on/contributed to the manuscript.

## Abbreviations

GTP: guanosine triphosphate
GDP: guanosine diphosphate
GMPCPP: guanosine-5’-[(α,β)-methyleno]triphosphate
GTPγS: guanosine-5’-O’[gamma-thio]triphosphate
+TIP: plus tip tracking protein
EB: end binding protein
FRAP: fluorescence recovery after photobleaching
MAP: microtubule associated protein
TIRF: total internal reflectance fluorescence

