## Supplemental Information for "Tau avoids the GTP cap at growing microtubule plus ends"

**Supplementary Methods:**

*Tubulin light scattering assay*

Polymerization of soluble tubulin was measured by monitoring the increase in scattered light at 340 nm, as previously described[28]. The tubulin was clarified as described above and buffer exchanged in BRB80 buffer (80 PIPES, 1 mM $MgCl_2$, 1 mM EGTA, and 1 mM DTT added immediately prior to use). For polymerization reactions, 20 μM tubulin was incubated with 2 μM tau for 2.5 minutes on ice prior to the addition of 1 mM GTP. Immediately after the addition of GTP, the reaction was transferred to a warmed cuvette and the reaction was monitored for 10 minutes at 37°C in a fluorometer (Fluorolog FL-1039/40, Horiba) with a photon counting module (SPEX DM302, Horiba) with both excitation and emission wavelengths set to 340 nm. Following polymerization, the samples were quickly returned to 4°C for 5 minutes; cold depolymerization is evidence that the proteins are not aggregated. The curves were normalized using Origin 2017.

*FCS instrument and data analysis*

All FCS measurements were performed on our home built instrument, as described previously [23,29]. Prior to entering the inverted Olympus 1X-71 Microscope (Olympus), the laser power was adjusted to ~ 5 μW (488 nm diode-pumped solid-state laser, Spectra-Physics) and focused into the sample via a 60x water immersion objective (Olympus). Fluorescence emission from the sample was collected through the objective, separated from excitation light by a Z488RDC long pass dichroic and a 500 nm long pass filter (Chroma). The filtered emission was focused the aperture of a 50 μm diameter optical fiber (OzOptics) coupled to an avalanche photodiode (Perkin-Elmer). A digital correlator (FLEX03LQ-12, Correlator.com) generated the autocorrelation curves.

Measurements were made in 8-chamber Nunc coverslips (Thermo-Fisher) passivated by incubation with (ethylene glycol)poly(L-lysine) (PEG-PLL) [28]. The labeled tau (15-25 nM) and tubulin (concentrations vary) were incubated in chambers for 5 minutes prior to measurement in phosphate buffer pH 7.4 (20 mM phosphate, 20 mM KCl, 1 mM $MgCl_2$, 0.5 mM EGTA, 1 mM DTT added immediately prior to use) at 20 °C. For GMPCPP-tubulin, buffer exchanged tubulin was incubated with 1mM GMPCPP for 5 mins prior to tau incubation. Multiple (20-40) 10 second autocorrelation curves were collected per sample and fit to a single-component 3D diffusion equation:

$$G(\tau) = \frac{1}{N\left(1+\frac{\tau}{\tau_D}\right)}\sqrt{\frac{1}{1+\frac{s^2\tau}{\tau_D}}} \qquad \text{Eq. S1}$$

where $G(\tau)$ is the autocorrelation function as a function of time ($\tau$), $\tau_D$ is the translational diffusion time of the labeled molecules and $N$ is the average number of fluorescent species. For our instrument, the ratio of the radial to axial dimensions of the focal volume ($s$) was determined to be 0.2 and consequently fixed for analysis.

The collected autocorrelation curves were analyzed as described previously (McKibben & Rhoades). Briefly, individual autocorrelation curves were fit with Eq. S1 and assessed the goodness of fit using least-squares $X^2 = [G(\tau)_{fit} - G(\tau)_{raw}]^2$ with a tolerance of $X^2 = 0.0001$ for a consecutive run of 75 ms. Infrequent and unusually large assemblies that pass this initial criterion still skew the data towards slower diffusion times. Although of potential interest in another context, these species do not represent the majority of the tau:tubulin complexes of interest here. These outliers were removed in an iterative fashion by testing the individual curves using an Anderson-Darling statistical test for either a lognormal or normal distribution. After testing the remaining autocorrelation curves were average together and fit according to Eq. S1 representing a single point. This was repeated in triplicate, and then numerically averaged.

<u>Figure S1:</u>

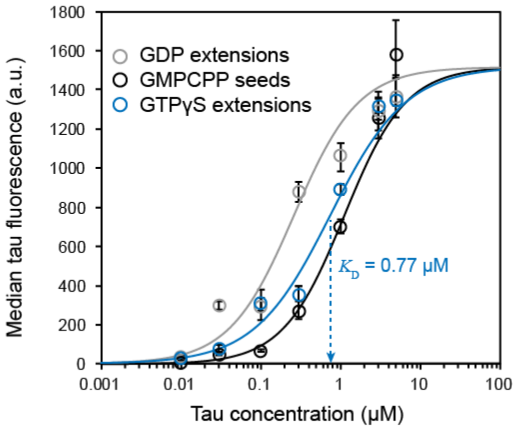


**Figure S1:** Median tau fluorescence on microtubule segments grown in the presence of different tubulin nucleotides *in vitro*, GDP (gray), GMPCPP (black), and GTPγS (blue). GDP and GMPCPP data is that shown in Figure 1C of the main text. Errorbars are ± 95% confidence interval.

### Figure S2:

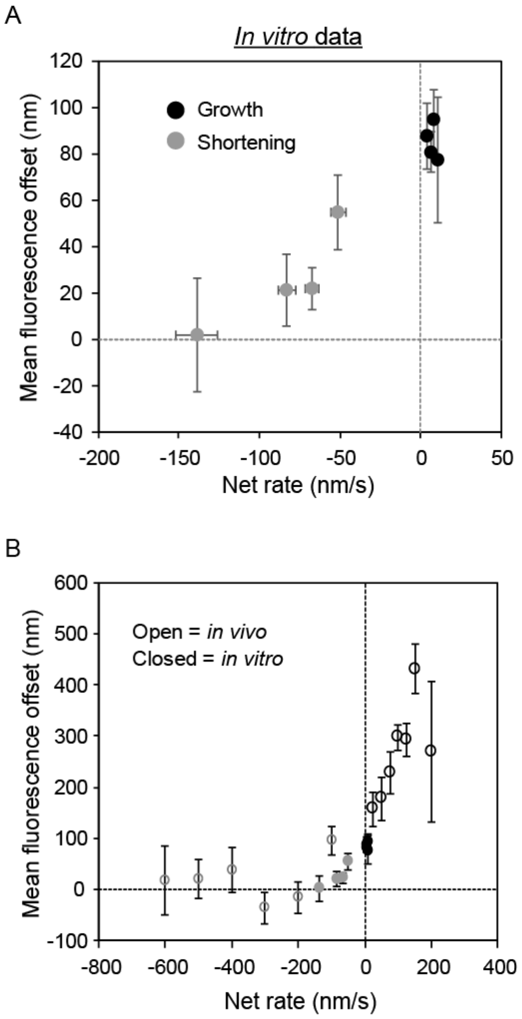


**Figure S2:** A) Mean tau fluorescence offset from the microtubule plus end as a function of the net rate of assembly *in vitro* only. Growing microtubules are in black while shortening microtubules are in gray. Errorbars are ± 95% confidence interval. B) Mean tau fluorescence offset from the microtubule plus end as a function of the net rate of assembly both *in vivo* and *in vitro*. Open circles are *in vivo* data and closed circles are *in vitro* data. Black indicates growing microtubules while gray indicates shortening microtubules. Errorbars are ± 95% confidence interval.

Figure S3:

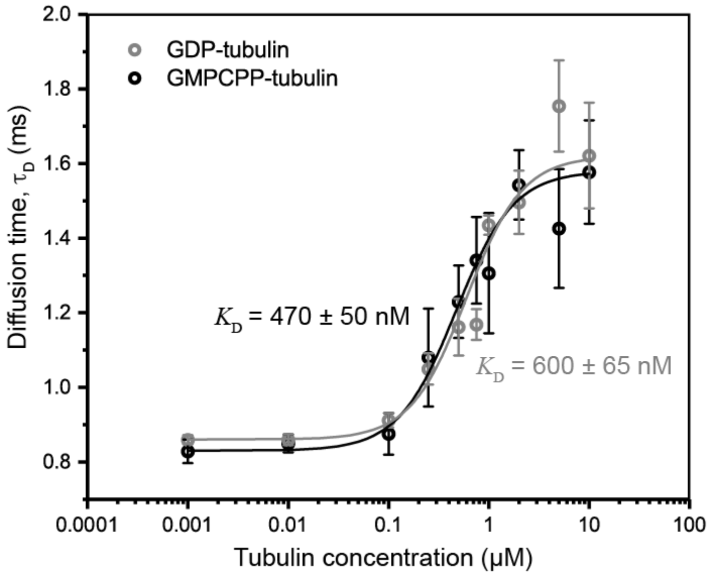


**Figure S3.** Tau does not exhibit a binding preference for GMPCPP-tubulin or GDP-tubulin in solution. Translational diffusion time measurements were made using fluorescence correlation spectroscopy (FCS) as previously described[23]. The increase in diffusion time as a function of tubulin concentration reflects binding of Alexa488-tau (20nM) to unlabeled tubulin rather than tubulin-tubulin oligomerization[29]. Measurements were carried out in phosphate buffer (pH 7.4) at 20ºC. Data are presented as mean ± 95% CI, $n \geq 3$ independent measurements. Curves indicate best-fit Hill function to all data points.

Figure S4:

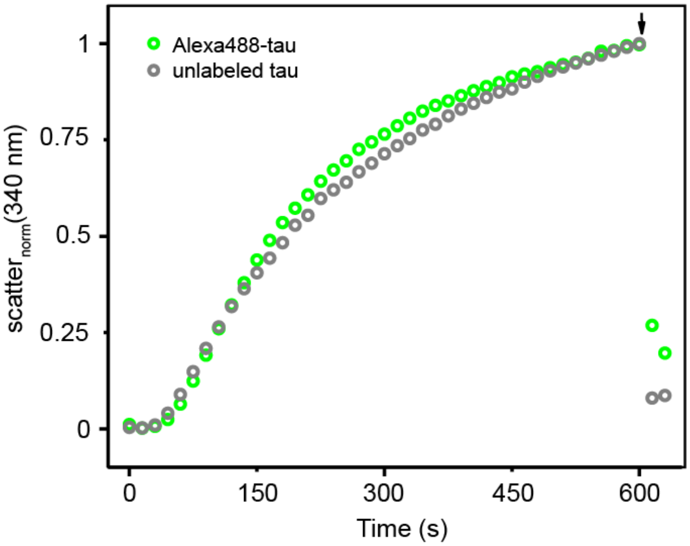


**Figure S4.** Labeled tau and unlabeled tau polymerize tubulin comparably. Normalized tubulin polymerization is shown as a function of time. Polymerization in the presence of Alexa488-tau is shown in green while that in the presence of unlabeled tau is shown in gray. Polymerization of soluble tubulin was measured by monitoring the increase in scattered light at 340 nm. For polymerization reactions, 20 µM tubulin was incubated with 2 µM tau for 2.5 minutes on ice prior to the addition of 1 mM GTP. The reaction was then transferred to a warm cuvette at 37ºC. The arrow indicates cold depolymerization at 4ºC.
